# Changes in gut microbiome composition drive fentanyl intake and striatal proteomic changes

**DOI:** 10.1101/2022.11.30.518531

**Authors:** Rebecca S. Hofford, Katherine R. Meckel, Weiwei Wang, Michelle Kim, Arthur Godino, TuKiet T. Lam, Drew D. Kiraly

**Author notes:** To whom correspondence should be addressed: Drew D. Kiraly, MD, PhD, Associate Professor Physiology, Pharmacology & Psychiatry, Wake Forest School of Medicine, Atrium Health Wake Forest Baptist, 115 S. Chestnut Street, Winston-Salem, NC 27104, Twitter: @kiralylab.

## Abstract

Opioid use disorder (OUD) is a public health crisis currently being exacerbated by increased rates of use and overdose of synthetic opioids, primarily fentanyl. Therefore, the identification of novel biomarkers and treatment strategies to reduce problematic fentanyl use and relapse to fentanyl taking is critical. In recent years, there has been a growing body of work demonstrating that the gut microbiome can serve as a potent modulator of the behavioral and transcriptional responses to both stimulants and opioids. Here, we advance this work to define how manipulations of the microbiome drive fentanyl intake and fentanyl seeking in a translationally relevant drug self-administration model. Additionally, we utilize global proteomic analysis of the nucleus accumbens following microbiome manipulation and fentanyl administration to define how microbiome status alters the functional proteomic landscape in this key limbic substructure. These findings establish clear relevance for gut-brain signaling in OUD, and lay foundations for further translational work in this space.

## Introduction

Opioid use disorder (OUD) is a public health problem leading to tremendous morbidity and mortality. Rates of opioid overdose have been increasing rapidly^1^ – an effect attributed to the increased presence of fentanyl in the drug supply^2^. Understanding the neurobiological factors driving fentanyl use and relapse is a critical step in identifying novel treatments to combat this epidemic.

The gut microbiome is a key regulator of health and homeostasis in diverse organ systems and disease processes. In recent years the understanding of the microbiome as a regulator of brain and behavior has increased markedly^3–5^. Recent work by our group and others has shown that alterations to the microbiome of rodents can affect cocaine^6,7^ and opioid reward and reinforcement^8,9^, can influence tolerance to opioids^10,11^, and can alter neuronal activation and transcriptional responses to opioids^8,12^. While this work was critical in establishing the role for gut-brain signaling in models of OUD, previous work has relied on transcriptional profiles and indirect measures of opioid reward as readouts for effects of the microbiome on brain and behavior.

Here, we utilize a translationally relevant fentanyl self-administration model to define effects of gut microbiome depletion on fentanyl self-administration and fentanyl-seeking after abstinence. These behavioral findings are coupled with detailed analysis of microbiome composition in relation to behavior, and quantitative proteomic analysis of the synaptic proteome of the nucleus accumbens (NAc). This work defines the gut microbiome as a mediator of fentanyl intake and seeking and identifies microbial communities, protein targets, and functional protein pathways that can be leveraged to reduce motivation to take and seek fentanyl.

## Materials & Methods

### Animals

Male Sprague-Dawley rats (Envigo, PND 80-90 at start of experiment) were pair-housed upon arrival to the colony. Rats remained pair-housed throughout the entire experiment. All rats were kept on a 12:12 hr reverse light cycle (lights on at 19:00) in a temperature and humidity-controlled vivarium. All procedures were approved by the IACUC at the Icahn School of Medicine at Mount Sinai and all experiments conformed to the standards provided in the “Guide for the Care and Use of Laboratory Animals”. Separate groups of rats were used for Experiments 1 and 2.

### Treatments

Cages were randomly assigned to H_2_O or Abx so both rats in a cage received the same drink type. Drink administration started 2 weeks before the start of self-administration. Abx mixture contained 0.5 mg/ml vancomycin (Chem-Impex International #00315), 2 mg/ml neomycin (Fisher Scientific #BP266925), 0.5 mg/ml bacitracin (Research Products International #B3200025), and 1.2 μg/ml pimaricin (Infodine Chemical #7681-93-8) in H_2_O. Control rats remained on H_2_O. Fentanyl hydrochloride was provided by the National Institute of Drug Abuse drug supply program and was dissolved in 0.9% saline.

### Fentanyl self-administration: General

Self-administration and the fentanyl-seeking test occurred in standard rat operant chambers with 2 fixed levers, two jewel lights above each lever, a house light, and a syringe pump (Med Associates, St Albans VT). All rats were implanted with a jugular catheter under ketamine/xylazine anesthesia (100/10 mg/kg). Jugular catheters (VABR1B/22, Instech, Plymouth Meeting PA) were inserted into each rat’s jugular vein, secured to the vein with silk thread, and threaded under their skin to exit their body via a back mount. Rats were allowed to recover individually until they regained consciousness before returning to pair housing. After a one-week recovery period, rats began self-administration as described below. Sessions occurred once daily and self-administration sessions lasted 3 hours during the rats’ dark phase. During self-administration, completion of the ratio requirements resulted in a 5.9 s infusion and the illumination of the light above the active lever for a 20 s time-out. Active and inactive lever placement was counterbalanced across all rats. Food was taken away from all rats 24 h before the start of acquisition and before the fentanyl seeking test; otherwise, rats were fed 18 g standard chow / rat during active self-administration and had *ad libitum* food access before self-administration started and during abstinence.

### Fentanyl self-administration Experiment 1: Increasing fixed ratio

Rats were trained to self-administer 2.5 microg/kg/infusion fentanyl or saline on a fixed-ratio 1 (FR1) schedule during acquisition which occurred once daily for 10 days. Following acquisition, fentanyl administering rats in both H_2_O and Abx groups were further divided into 2 groups-half of the rats underwent an increasing fixed ratio assessment for 6 days (referred to as H_2_O-IncFR & Abx-IncFR throughout). This progressed as FR2 for 2 days, FR3 for 2 days, and FR5 for 2 days; the other half of the rats continued maintenance at an FR1 for 6 days (referred to as H_2_O-FR1 & Abx-FR1). Saline-administering rats underwent increasing FR as described above (H_2_O-Sal & Abx-Sal). Immediately after the increasing FR / maintenance phase, all rats had 2 consecutive days of progressive ratio. The response requirement for each successive infusion during progressive ratio followed the formula: response ratio = [5e^(injection number x 0.2)] – 5^13^. After progressive ratio, all rats underwent 2 days FR1 before 20 days of home cage abstinence. Finally, after abstinence, rats were tested on a combined context + cue fentanyl seeking test where rats were placed in the operant boxes with both levers extended. Lever presses were recorded but had no programmed consequence for 30 minutes. At 30 minutes, the light above the active lever was illuminated for 20 s. After this, every lever press on the previously active lever resulted in illumination of the previously active lever light. Inactive lever presses continued to have no consequence. This continued for another 30 mins. Twenty-four hours after their seeking test, rats were euthanized; nucleus accumbens (NAc) tissue and cecal contents were collected and flash-frozen on dry ice. Rats were dropped from the study if they lost catheter patency or did not administer more than 10 infusions per day over the last 3 days of acquisition.

### Fentanyl self-administration Experiment 2: Dose-response

Rats from H_2_O and Abx groups were trained to self-administer 2.5 microg/kg/infusion fentanyl on an FR1 once daily for 10 days. Immediately after their last acquisition session, rats were allowed to self-administer fentanyl at the following doses: 0, 0.025, 0.25, 0.79, 7.9, and 25 microg/kg/infusion. Rats received 2 consecutive days of every dose, but dose order across rats was randomized. Rats were dropped from the study if they lost catheter patency or did not administer more than 10 infusions per day over the last 3 days of acquisition.

### Statistics for behavioral experiments

To minimize excessive comparisons, H_2_O and Abx groups were compared directly to each other only if they were administering the same drug (fentanyl or saline) and they underwent the same cycle of self-administration phases (e.g. Experiment 1 increasing FR or Experiment 1 maintenance). Active and inactive lever presses during acquisition, increasing fixed ratio, maintenance, and FR1 after progressive ratio were analyzed separately using linear mixed effects with drink type as a fixed between-subjects factor and session or dose as a fixed within-subjects factor. Active and inactive lever presses during the dose-response were analyzed with a two-way ANOVA. Progressive ratio breakpoints and lever presses during the fentanyl-seeking test were analyzed using unpaired t-tests between H_2_O and Abx for each set of rats administering the same drug and undergoing the same cycle of self-administration phases. In the presence of significant interactions, differences between H_2_O and Abx at all sessions or doses were analyzed with Tukey’s LSD. Greenhouse-Geiser corrections were applied and statistical outliers were removed when appropriate.

### 16S sequencing

Microbial DNA was isolated from the cecal contents of rats from Experiment 1 using Qiagen DNeasy PowerSoil Pro kit per kit instructions. DNA concentration was determined with a NanoDrop1000. PCR amplification was achieved using primers (341F/805R) targeting the V3 and V4 region of the 16S rRNA region of the bacterial genome and was sequenced on an Illumina NovaSeq (2 × 250 bp paired-end). Amplicons were chimera filtered, dereplicated, and paired-ends were merged using Divisive Amplicon Denoising Algorithm 2 (DADA2). These features were used to determine observed taxonomic units (OTU), defined as sequences with ≥ 97% similarity. OTU counts were used to determine the Shannon index of alpha diversity, the Simpson index, and principle coordinates analysis plots were generated using the Unifrac distance as an assessment of beta diversity using QIIME2 software. OTUs were identified by comparing their genetic sequences to reference bacterial genomes using SILVA (Release 132) with confidence set at 0.7. Fold change phyla abundance from H_2_O Sal were analyzed using independent t-tests. Genera abundance was correlated with average number of infusions earned during the last 2 days of increasing FR / maintenance.

### Sample Preparation for LC–MS/MS & Data-Independent Acquisition

Nucleus accumbens tissue was processed for protein isolation, and data-independent analysis tandem mass spectrometry largely as previously described^14^, and as described in detail in the **Supplementary Methods**.

### Pathway analysis and visualization of protein networks

Proteins were excluded from analysis if they were not detected in > 50% of all samples irrespective of treatment. Pairwise comparisons of the Log_10_ median intensity of every remaining protein and protein group were made using Scaffold DIA proteomics analysis software (http://www.proteomesoftware.com/products/dia/). Technical replicates were treated as independent samples and proteins were considered significantly differentially regulated when FDR corrected *p* < 0.1. All groups were compared to H_2_O-Sal to allow for inferences across comparisons. Significantly upregulated and downregulated proteins were separately uploaded into the open source pathway analysis software package G:Profiler^15^ (https://biit.cs.ut.ee/gprofiler/gost) to identify significantly enriched Gene Ontologies (GO) and Kyoto Encyclopedia of Genes and Genomes (KEGG) pathways. Enrichr^16,17^ (https://maayanlab.cloud/Enrichr/) was used to identify upstream predicted transcription factors using the “ENCODE and CheA consensus transcription factors from Chip-X” list using an FDR corrected *p* < 0.05. Additionally, all significantly regulated proteins were uploaded to Ingenuity Pathway Analysis (IPA) for analysis of pathway directionality between comparisons using an FDR corrected *p* < 0.05. A list of shared upregulated proteins between fentanyl self-administration groups that excluded proteins upregulated in Abx Sal was generated using DeepVenn^18^ (http://www.deepvenn.com/); these proteins were separately uploaded into the STRING database^19^ (https://string-db.org/). Cytoscape with STRING add-in was used for visualization of protein-protein interaction within the IPA pathway “synaptogenesis signaling pathway”. Given that G:Profiler, Enrichr, STRING, and IPA use gene names for identifying pathways, all protein names were first converted to gene names prior to analysis using the Uniprot database (https://www.uniprot.org/id-mapping). Portions of figures 1, 2, and 3 were made with Biorender.com.

**Fig. 1.**
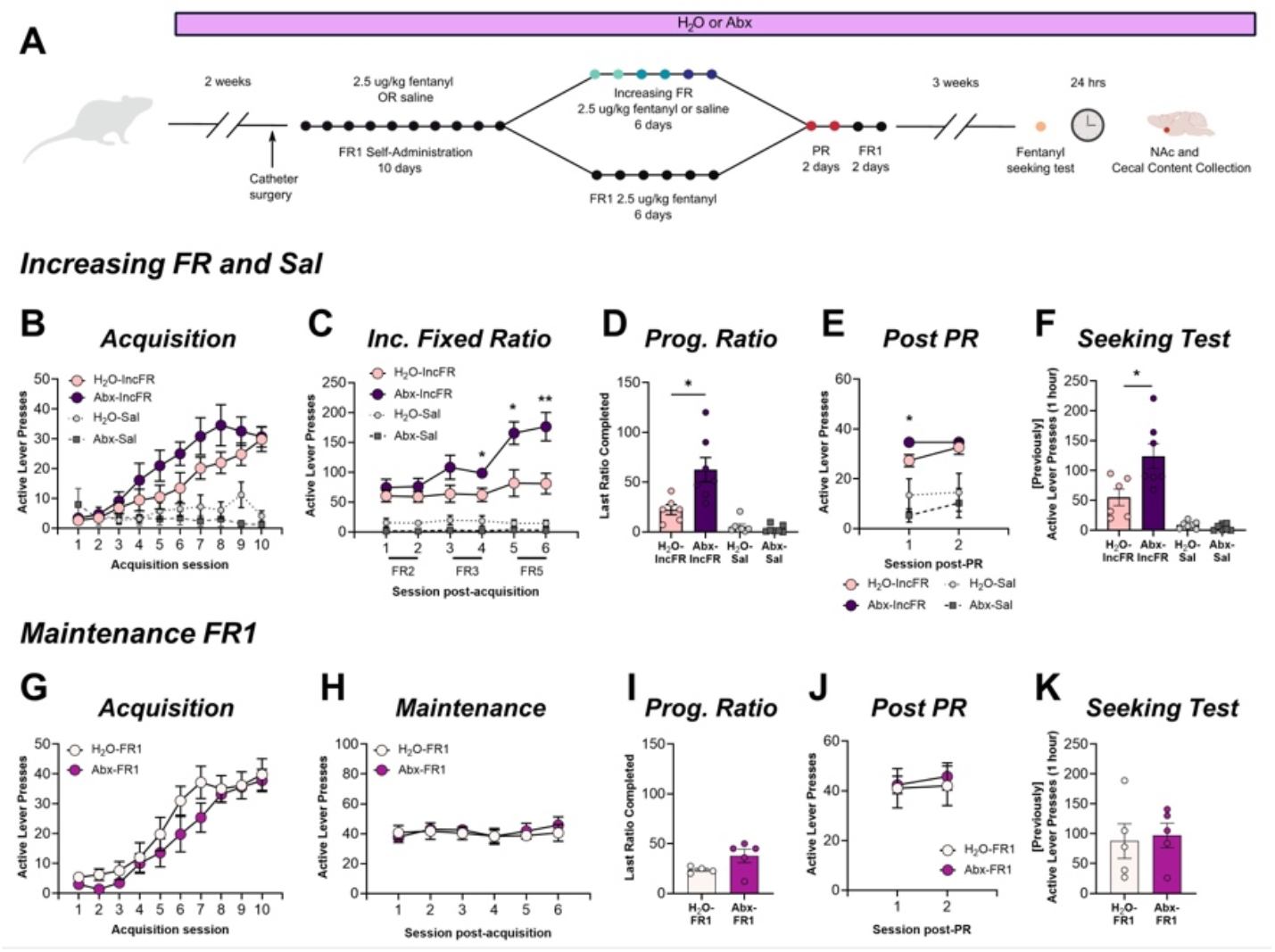
Microbiome depletion increases motivation to self-administer fentanyl. **(A)** Experimental timeline for Experiment 1. (**B)** H_2_O-IncFR and Abx-IncFR rats acquired FR1 fentanyl administration at equal rates. **(c)** At increasing FR requirements Abx caused enhanced responding. **(D)** Abx-IncFR rats had higher breakpoints on a progressive ratio task. **(E)** Return to FR1 normalized responding between groups over two sessions. **(F)** Abx treatment increased fentanyl-seeking after withdrawal in Abx-IncFR rats. **(G-K)** As a control, another group of rats was maintained on FR1 responding throughout (H_2_O-FR1, Abx-FR1), and there were no differences in acquisition of self-administration, maintenance of self-administration, progressive ratio, post-PR FR1 responding, or fentanyl-seeking. Data presented as means ± SEM. * *p* < 0.05, ** *p* < 0.01. Full statistical results for active and inactive lever presses are included in Supplemental tables 1-2.

**Fig. 2.**
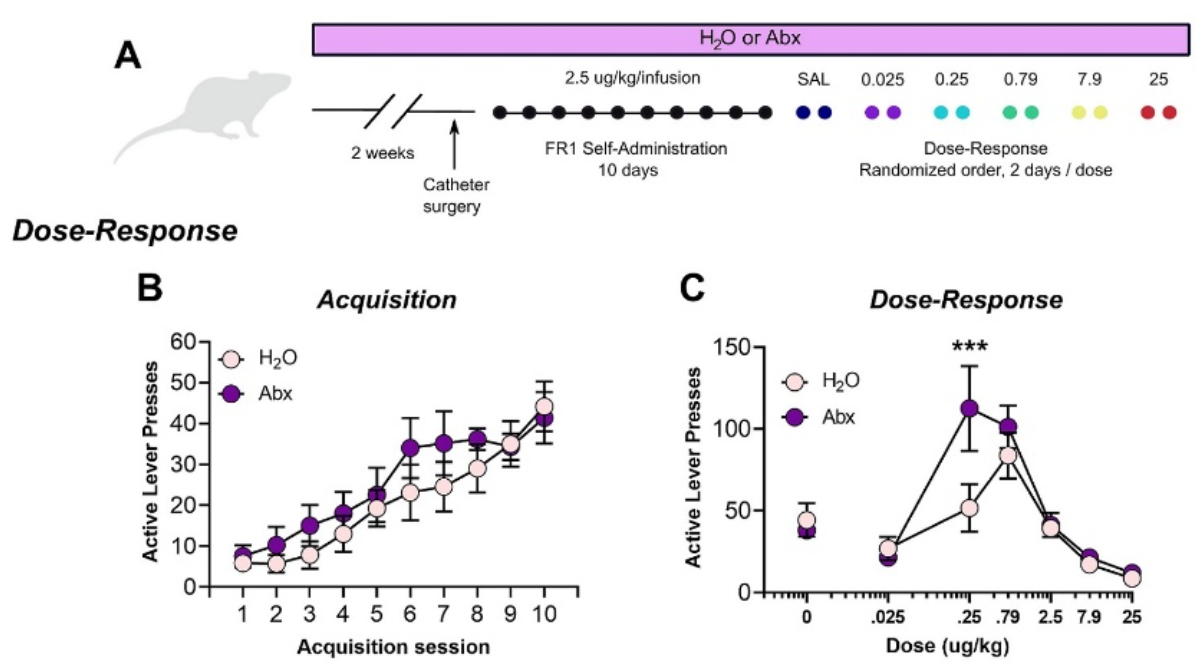
Microbiome depletion shifts the dose-response curve. **(A)** Experimental timeline for Experiment 2. **(B)** Abx did not affect acquisition of self-administration. **(C)** However, there was a significant interaction between dose and Abx treatment, post hoc test indicated a significant difference between groups at the 0.25 microg/kg/infusion dose. Data presented as means ± SEM. *** *p* < 0.001. Full statistical results for active and inactive lever presses are included in Supplemental tables 1-2.

**Fig. 3.**
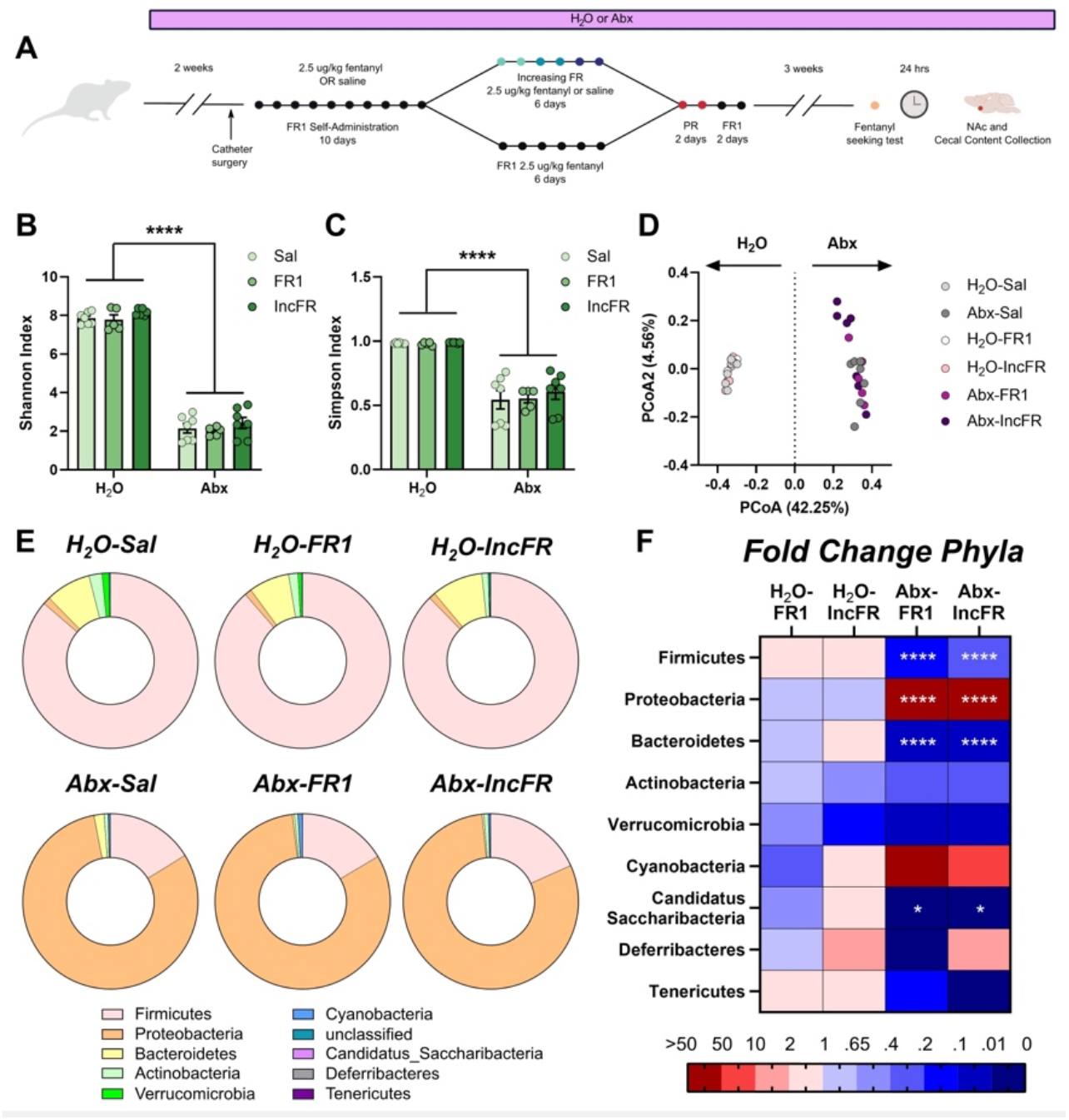
Abx, but not fentanyl, alter the microbiome. **(A)** Experimental timeline for Experiment 1 and microbiome collection. Abx reduced microbiome diversity as measured by the Shannon **(B)** and Simpson **(C)** indices. **(D)** The microbiomes of H_2_O and Abx rats differed markedly. Microbial composition was driven primarily by Abx treatment with no observed effect of fentanyl self-administration compared to saline controls. **(E)** Donut plots of bacterial phyla abundance in all six groups of rats. **(F)** Heatmap of phyla abundance as fold change from H_2_O Sal in fentanyl-administering groups of rats. * *p* < 0.05, *** *p* < 0.001, *** *p* < 0.0001. Data presented as means ± SEM.

## Results

### Microbiome knockdown enhances motivation for fentanyl and induces a leftward and upward shift in the dose-response curve

To investigate the consequence of microbiome knockdown on fentanyl intake and motivation, rats in Experiment 1 were treated with antibiotics (Abx) to reduce the microbiome or maintained on control water (**Fig. 1A**). As expected, microbiome knockdown did not alter saline intake (session: *F*_(1.83, 22.01)_=1.01, *p*=0.38; drink: *F*_(1,12)_=0.57, *p*=0.47, interaction: *F*_(9,108)_=2.73, *p*=0.007, but no significant pairwise comparison were identified) and did not affect acquisition of fentanyl self-administration when trained at an FR1 (main effect of session only *F*_(1.62,17.66)_=29.9, *p*<0.0001; drink: *F*_(1,11)_ = 2.07, *p* = 0.18; interaction: *F*_(9, 98)_=1.361, *p* = 0.22). However, when rats self-administered at higher fixed ratios, Abx-IncFR rats increased effort to obtain fentanyl as it became harder to earn (**Fig. 1C**, session: *F*_(2.06,22.19)_=25.51, *p*<0.0001; drink: *F*_(1,11)_=5.39, *p*=0.04; and interaction: *F*_(5,54)_=9.94, *p*<0.0001), and had higher breakpoints on a progressive ratio (PR) task (**Fig. 1D**, *t*_(11)_=2.88, *p*=0.02). Since the fixed ratio requirements increased over time (FR2/3/5) and opioid tolerance can change over time in a microbiome-dependent manner^10,11^, a separate group of fentanyl-administering rats were kept at FR1 (referred to as H_2_O-FR1 and Abx-FR1) and were run concurrently (**Fig. 1A**, bottom). When maintained at FR1, microbiome knockdown had no effect on acquisition of fentanyl self-administration (**Fig. 1G**, session: *F*_(2.02,15.71)_=47.33, *p*<0.0001; drink: *F*_(1,8)_=1.18, *p*=0.31; interaction: *F*_(9,70)_=0.98, *p*=0.47), maintenance of FR1 responding over time (**Fig. 1H**, session: *F*_(1.98,15.42)_=0.57, *p*=0.57; drink: *F*_(1,8)_=0.11, *p*=0.74; and interaction: *F*_(5,39)_=0.36, *p*=0.87), or with PR testing (**Fig. 1I**, *t*_(7)_=1.77, *p*=0.53). Thus, Abx-induced enhancement in responding for fentanyl during the increasing FR phase represents an increase in motivation for fentanyl instead of altered tolerance or a shifting microbiome.

Following PR testing, all rats were restabilized on FR1 responding (**Figs. 1E and 1J**). After 20 days of abstinence, Abx-IncFR rats showed markedly higher levels of fentanyl-seeking during a context and cue fentanyl-seeking task (**Fig. 1F**, *t*_(7)_=2.66, *p*=0.02), but this was not seen with Abx-FR1 rats (**Fig. 1K**, *t*_(8)_=0.27, *p*=0.79).

Since microbiome knockdown led to differences in motivation to administer only at higher FRs, we considered the possibility that microbiome disruption causes a shift in the dose-response curve. To test this, Experiment 2 utilized a separate group of rats that were treated and trained to administer fentanyl as described above (**Fig. 2A**). Like previously observed, Abx did not alter the rate of acquisition (**Fig. 2B**, session: *F*_(1.65,19.05)_=26.85, *p*<0.0001; drink: *F*_(1,12)_=0.87, *p*=0.37; interaction: *F*_(9,104)_=0.75, *p*=0.63). However, there was a significant interaction between treatment and dose during assessment of their dose-response curves (**Fig. 2C**, interaction: *F*_(5,66)_=2.61, *p*=0.03; dose: *F*_(5,66)_=21.09, *p*<0.0001; drink: *F*_(1,66)_=4.99, *p*=0.03), demonstrating a leftward and upward shift of the curve after microbiome knockdown, and a significant post-hoc difference between groups at the 0.25 microg/kg dose (*p*=0.0008). Full active lever and inactive lever statistics included in **Supplemental tables 1-2**.

### Fentanyl intake during increasing FR and maintenance is negatively correlated with several bacterial genera

We next performed detailed analysis of changes to the microbiome caused by Abx and fentanyl self-administration using 16S sequencing of cecal contents from rats in Experiment 1 (**Fig. 3A**). As expected, Abx but not fentanyl exposure, drastically reduced indices of alpha diversity (**Figs. 3B-C**, Shannon index drink: *F*_(1, 31)_ = 1123.0, *p* < 0.0001; self-administration paradigm: *F*_(2, 31)_ = 1.97, *p* = 0.16; interaction: *F*_(2, 31)_ = 0.01229, *p* = 0.9878; Simpson index drink: *F*_(1, 31)_ = 129.3, *p* < 0.0001; self-administration paradigm: *F*_(2, 31)_ = 0.3608, *p* = 0.7; interaction: *F*_(2, 31)_ = 0.2261, *p* = 0.80). Additionally, as observed in our prior study utilizing morphine^8^, we found that opioid history did not significantly alter the microbiome - there was no separation of fentanyl and saline samples assessed using a Unifrac dissimilarity matrix (**Fig. 3D**). Visualization of phyla abundance across groups indicated that relative abundance of Firmicutes, Bacteroidetes, and Candidatus Saccharibacteria were reduced and levels of Proteobacteria were expanded after Abx (**Figs. 2E-F**, full statistics on phyla and genus abundance included in **Supplemental tables 3-4**).

Next, we examined the relationship between fentanyl intake and genus abundance within our H_2_O groups (H_2_O-FR1 and H_2_O-IncFR). Abundance of *Ruminococcus, Butyricicoccus, Lachnospiracae_unclassified*, and *Anaerotignum* negatively correlated with fentanyl intake during the last 2 days of fentanyl increasing FR or maintenance (**Figs. 4A-D**). Interestingly, the abundance of these four genera was markedly diminished by Abx treatment (**Figs. 4A-D, insets**), suggesting that low levels of these bacteria could enhance motivation for and sensitivity to the reinforcing properties of fentanyl observed here after microbiome knockdown. Correlation statistics included in **Supplemental table 5**.

**Fig. 4.**
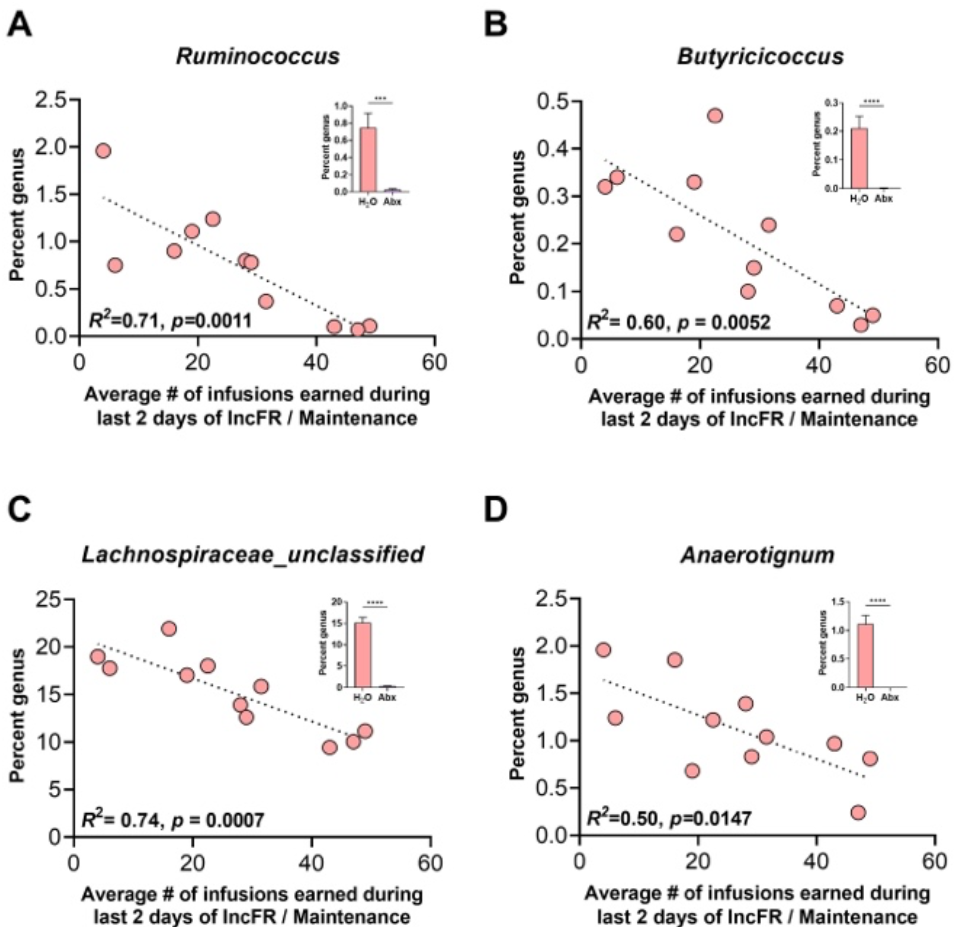
Several genera correlate with fentanyl intake in rats with an intact microbiome. Correlation plots of H_2_O fentanyl rats’ fentanyl intake with relative abundance of **(A)** *Ruminococcus*, **(B)** *Butyricicoccus*, **(C)** *Lachnospiraceae_unclassified*, and **(D)** *Anaerotignum*. Insets: percent abundance of each genus in H_2_O and Abx groups, collapsed across administration paradigm (FR1 and IncFR combined). * *p* < 0.05, *** *p* < 0.001, *** *p* < 0.0001. Data presented as means ± SEM.

### Microbiome knockdown alters global protein expression after a fentanyl-seeking task

We previously found that microbiome knockdown altered the effect of repeated morphine on transcriptional control in the NAc^8^. Knowing that microbiome knockdown has such a robust effect on transcriptional responses to opioids and that exposure to drug-related cues after a period of abstinence contributes to relapse^20^, the current study examined how microbiome depletion alters the proteome in the NAc at a critical time point after a fentanyl-seeking test (**Fig. 5A**). Both Abx treatment and fentanyl self-administration affected the proteomic landscape but rats in the Abx-IncFR group had the most differentially expressed proteins compared to H_2_O-Sal controls (**Fig. 5B**, full lists of differentially regulated proteins included as **Supplemental table 6**). Since behavioral differences were strongest between H_2_O- and Abx-IncFR groups, pathway analysis focused on protein expression differences between these groups. A subset of pathways were uniquely predicted in either H_2_O or Abx-IncFR including pathways ‘regulation of presynaptic cytosolic calcium’ in H_2_O-IncFR and ‘small molecule binding’ in Abx-IncFR. (**Supplemental table 7** and **Supplemental fig. 3**). However, most of the top terms associated with substance use disorders (SUD) were present in both groups (**Fig. 5C**), including the predicted upregulated pathways: ‘amphetamine addiction’, ‘cocaine addiction’, and ‘dopaminergic synapse’.

**Fig. 5.**
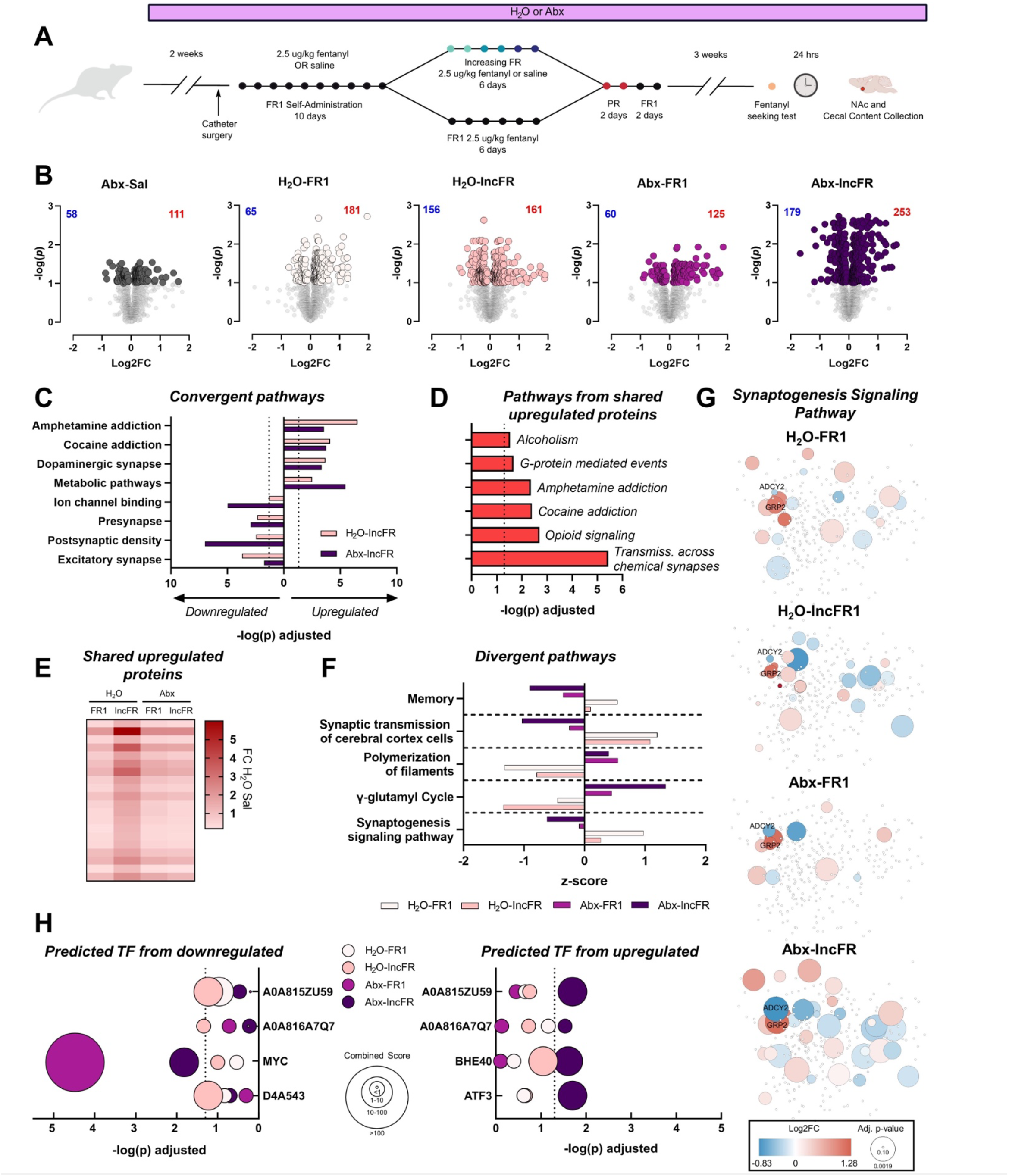
Microbiome depletion alters the nucleus accumbens proteome. **(A)** Experimental timeline for Experiment 1 and NAc collection. **(B)** Volcano plots depicting protein expression changes in rats compared to H_2_O Sal. Colored points are significantly regulated proteins (FDR-corrected *p* < 0.1). **(C)** Select pathways that are predicted to be regulated in the same direction in H_2_O-IncFR and Abx-IncFR groups. Dotted line at 1.3 indicates significance (FDR-corrected *p* < 0.05). **(D)** Predicted pathways from proteins upregulated in all fentanyl-administering groups but not present in Abx-Sal. **(E)** Heatmap of fold change protein expression of overlapping upregulated proteins in fentanyl-administering rats. **(F)** Select pathways that are oppositely regulated in H_2_O and Abx groups. Note X-axis is z-score; all pathways in all groups are significant (FDR-corrected *p* < 0.05). **(G)** Cloud diagrams of proteins in the “synaptogenesis signaling pathway”. Each diagram presents the same configuration of proteins with differences between groups indicated by the colored protein nodes. **(H)** Transcription factors predicted to be upstream from downregulated proteins (*left*) and upregulated proteins (*right*).

Given the overlap in predicted pathways, we performed pathway analysis on shared proteins that were up-or downregulated in all fentanyl administering groups and were unchanged in Abx-Sal. There were 3 shared downregulated proteins and 20 shared upregulated proteins (**Fig. 5E**). Notably, several proteins that have been implicated in the actions of other drugs of abuse^21,22^ were upregulated in all fentanyl groups such as PDYN, which had the highest fold change difference in every comparison, and SC6A3, commonly known as the dopamine transporter. No pathways were predicted from downregulated proteins, but several pathways were predicted from upregulated proteins (**Supplemental table 8**) including the SUD-related terms: ‘alcoholism’, ‘amphetamine addiction’, and ‘opioid signaling’ (**Fig. 5D**).

Using Ingenuity Pathway Analysis, which provides directional z-scores from total lists of differentially regulated proteins, several pathways were significantly predicted to be regulated in opposite directions in H_2_O rats versus Abx rats. Among them, ‘memory’ and ‘synaptogenesis signaling pathway’ were predicted to be downregulated in Abx groups and upregulated in H_2_O groups (**Fig. 5F**, full list of IPA terms as **Supplemental tables 9-10**). These pathways were strongly regulated by microbiome status, indicating that these pathways might be important in driving the behavioral differences observed in the current study. To visualize the extent of this discrepancy, cloud diagrams depicting proteins within the synaptogenesis signaling pathway were generated (**Fig. 5G**, full list of proteins in **Supplemental table 11**).

Finally, upregulated and downregulated protein lists were analyzed using Enrichr database to identify potential transcription factors that might be regulating the changes in protein expression observed. Only A0A815ZU59 and A0A816A7Q7 (gene names *Zmiz1* and *Yy1*, respectively) were significantly predicted to be oppositely regulated between H_2_O-IncFR and Abx-IncFR (**Fig. 5H**, full list of predicted transcription factors as **Supplemental table 12**). Additional transcription factors were differentially predicted in some groups; notably, MYC was highly downregulated in all Abx groups but not in any H_2_O groups, suggesting that activity of MYC could be modulated directly by the microbiome and could be affecting motivation and fentanyl-seeking.

## Discussion

These studies provide evidence that the microbiome influences opioid reinforcement in a translationally relevant model of opioid use. By utilizing self-administration, we show that microbiome knockdown increases motivation for fentanyl, enhances sensitivity of the reinforcing properties of fentanyl, and increases fentanyl-seeking (**Figs. 1-2**). Previously, we observed that mice with a depleted microbiome have reduced morphine conditioned place preference at higher doses of morphine^8^. While there are several experimental differences between the current work and the previous report (e.g. species, opioid, and behavioral test), we think the morphine CPP results and the self-administration data can generally be explained by the same behavioral mechanism. As observed in Experiment 2, rats show enhanced sensitivity to the rewarding effects of fentanyl-demonstrated by the leftward shift in the dose-response curve. While involving different types of learning (instrumental versus associative), both lever pressing for infusions during self-administration and conditioned place preference^23^ are dose-dependent and the dependent variable-dose relationship is shaped as an inverted U. Thus, a *reduction* in high dose morphine place preference could be explained by a leftward shift in the CPP dose-response curve. In line with our results, other laboratories have described a similar enhancement of lower-dose morphine’s effectiveness in tests of antinociception after microbiome knockdown^11^.

As observed previously, Abx drastically reduced levels of bacterial diversity and significantly shifted the microbiome. Notably, alterations in multiple antibiotic-sensitive genera negatively correspond with fentanyl intake (**Fig. 4**). Four genera (*Ruminococcus, Butyricicoccus, Lachnospiracae_unclassified*, and *Anaerotignum*) were significantly reduced in Abx-treated rats, suggesting that lower levels of these bacteria (whether induced by oral Abx or occurring naturally in the H_2_O-treated rats) could be responsible for greater fentanyl intake at higher fixed ratios. While the relationship between genus abundance and fentanyl intake is a correlation and directionality of effect cannot be determined with certainty, it is unlikely that greater fentanyl intake caused a reduction in genera abundance. As we have observed previously in mice^8^, opioid exposure does not significant influence microbiome diversity. Similarly, fentanyl exposure did not affect microbiome similarity as assessed by Unifrac dissimilarity, a sensitive metric of microbiome diversity between samples. Interestingly, *Ruminococcus, Butyricicoccus*, and *Lachnospiraceae_unclassified* are all within the Clostridia class of bacteria; species within this class are producers of short-chain fatty acids (SCFA)^24^. We have previously shown that SCFA supplementation can reverse the effects of Abx on morphine CPP^8^, indicating a possible role for SCFA in altered opioid self-administration as well.

Finally, we performed proteomic analysis on NAc samples from rats that self-administered fentanyl or saline in Experiment 1. With the inclusion of saline controls (H_2_O-Sal and Abx-Sal) as well as H_2_O and Abx groups that self-administered the same amount of fentanyl (H_2_O-FR1 and Abx-FR1) and H_2_O and Abx groups that differed in their fentanyl intake (H_2_O-IncFR and Abx-IncFR), the design of this experiment allowed us to separate protein expression differences driven by Abx alone from protein expression alterations that were driven by differences in fentanyl administration secondary to microbiome depletion. With this in mind, we identified several pathways that were significantly regulated in different directions by H_2_O and Abx regardless of self-administration paradigm, suggesting that processes related to these pathways such as memory or synaptogenesis signaling drive the behavioral effects of Abx (**Fig. 5**). Additionally, we identified several proteins that were upregulated in all fentanyl-administering groups regardless of microbiome status including PDYN and the dopamine transporter-proteins not previously linked to fentanyl but implicated in psychostimulant relapse. Finally, we identified MYC as a potential transcription factor of interest-its activity is predicted to be significantly down within Abx groups. This set of studies defines gut-brain signaling that can drive fentanyl intake and relapse and lays the groundwork for future translational studies that can be harnessed to curb the OUD epidemic.

## Acknowledgements

Fentanyl hydrochloride was provided by NIDA drug supply. We also thank Florine Collin from the Keck MS & Proteomics Resource at Yale School of Medicine for assistance with preparing the proteomics samples. Schematics in all figures were prepared using BioRender.com with full permission to publish.

## Author Contributions

R.S.H. and D.D.K. designed the experiments. R.S.H., K.R.M., M.K., A.G., W.W. & T.T.L. performed the experiments. R.S.H., W.W., T.T.L. & D.D.K. performed data analysis. R.S.H. & D.D.K. wrote the manuscript. All authors critically edited and approved the final version of the manuscript.

## Supplemental Figures

**Supplemental Fig. 1:**
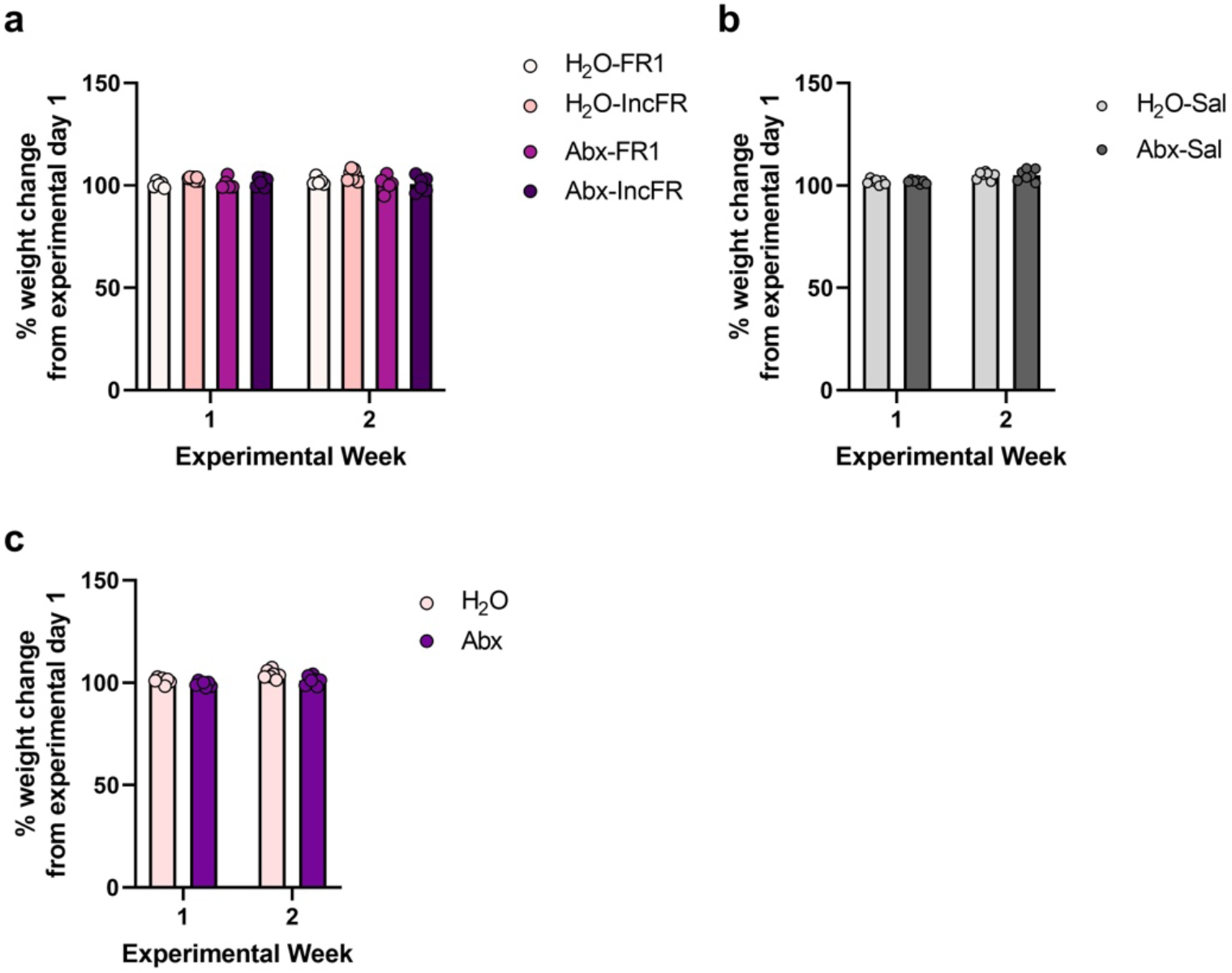
Abx alone does not affect rat body weight gain. For Experiment 1, there were no significant effects on body weight change between **(a)** H_2_O-FR1, H_2_O-IncFR, Abx-FR1, and Abx-IncFR (mixed effects, group: *F*_*(3, 19)*_ = 3.038, *p* = 0.0543; experimental week: *F*_*(1, 19)*_ = 1.359, *p* = 0.2582; interaction: *F*_*(3, 19)*_ = 2.475, *p* = 0.0927) or (**b)** between H_2_O-Sal and Abx-Sal rats (mixed effects, group: *F*_*(1, 12)*_ = 0.004688, *p* = 0.9465; experimental week: *F*_*(1, 12)*_ *=* 21.33, \*\*\**p* = 0.0006; interaction: *F*_*(1, 12)*_ = 0.002415, *p* = 0.9616). **(c)** For Experiment 2, Abx treatment minimally reduced body weight change (mixed effects, group: *F*_*(1, 12)*_ *=* 7.245, \**p* < 0.0196; experimental week: *F*_*(1, 12)*_ *=* 41.85, \*\*\*\**p* < 0.0001; interaction: *F*_*(1, 12)*_ = 2.335, *p* = 0.1524). However, average percent weight change only differed by 3 percentage points between H_2_O and Abx at both time points (Week 1: H_2_O average percent weight change = 101.06%, average Abx weight change = 99.33%; Week 2: average H_2_O weight change = 104.02%, average Abx weight change = 101.16%). Data presented as means ± SEM. * *p* < 0.05, ** *p* < 0.01, *** *p* < 0.001, **** *p* < 0.0001.

**Supplemental Fig. 2:**
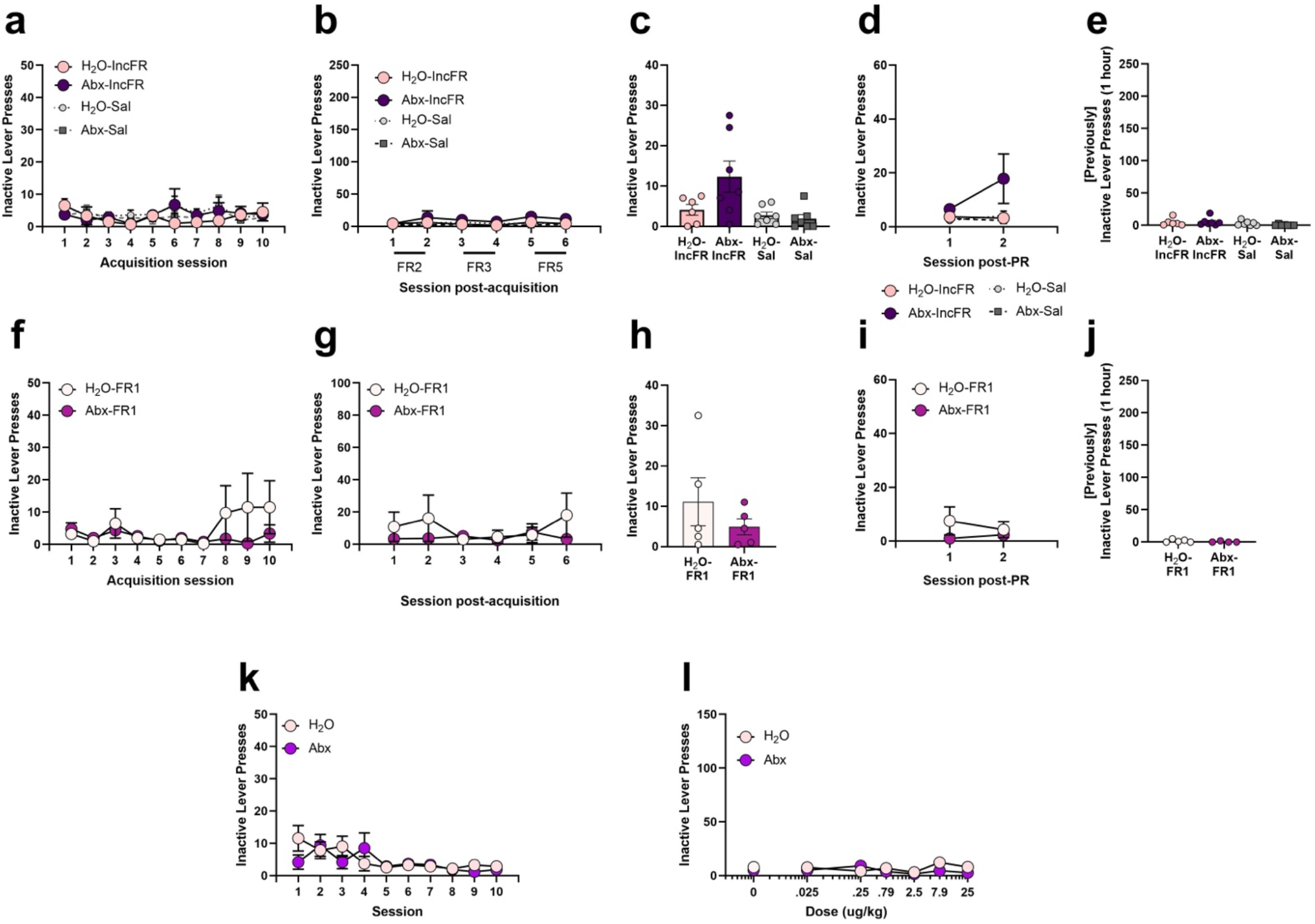
Microbiome knockdown does not affect inactive lever press responding. In Experiment 1, Abx did not affect inactive lever presses during **(a)** acquisition, **(b)** increasing fixed ratio, **(c)** progressive ratio, **(d)** post-PR FR1 responding, or **(e)** during a fentanyl-seeking test; no differences were seen between H_2_O-IncFR and Abx-IncFR groups or H_2_O-Sal and Abx-Sal groups. Additionally, there were no differences between H_2_O-FR1 and Abx-FR1 during **(f)** acquisition, **(g)** maintenance, **(h)** progressive ratio, **(i)** post-PR FR1 responding, or **(j)** during fentanyl-seeking. For Experiment 2, there was an interaction between Abx and session during **(k)** acquisition (*F*_*(9, 99)*_ = 1.98, *p* = 0.0495) but post hoc analysis found no difference between H_2_O and Abx on any single day; additionally inactive lever presses were lower than 5 for both groups over the last 5 sessions.. Abx did not influence inactive lever pressing during assessment of the **(l)** doseresponse. Full statistics for inactive lever presses are listed in **Extended Data Table S2**. * *p* < 0.05, *** *p* < 0.001, *** *p* < 0.0001. Data presented as means ± SEM.

**Supplemental Fig. 3:**
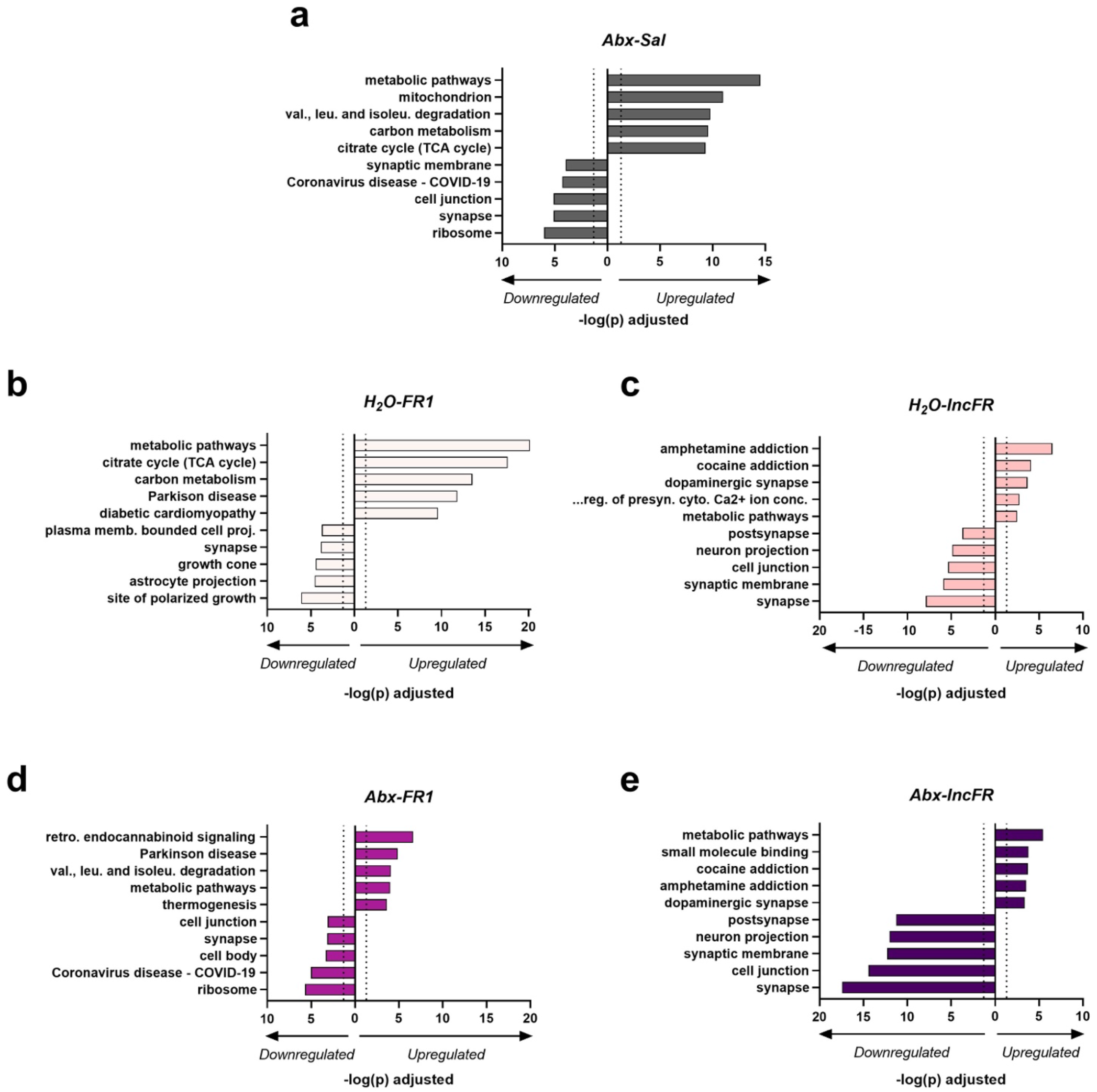
Top gene ontology pathways. Most significantly regulated gene ontology pathways in **(a)** Abx-Sal, **(b)** H_2_O-FR1, **(c)** H_2_O-IncFR, **(d)** Abx-FR1, and **(e)** Abx-IncFR groups compared to H_2_O-Sal. Dotted lines at 1.3 indicate significance (FDR-corrected *p* < 0.05).

